## Supplementary Information for "Telomerase RNA structural heterogeneity in living human cells detected by DMS-MaPseq"


| | |
|---|---|
| Supplementary Table S1. | Oligonucleotides used in the study. |
| Supplementary Figure S1. | Replicate DREEM-deconvoluted DMS profiles of hTR t/PK. |
| Supplementary Figure S2. | Quantitative evaluation of DMS-modification data concordance with predicted RNA secondary structure models. |
| Supplementary Figure S3. | Replicate DREEM-deconvoluted DMS profiles of hTR CR4/5. |
| Supplementary Figure S4. | Unguided RNAstructure predictions of hTR CR4/5. |
| Supplementary Figure S5. | DREEM-deconvoluted DMS profiles of the CR4/5 domain in BJ Fibroblasts. |
| Supplementary Figure S6. | Comparison of DREEM-deconvolution results across different cell types. |
| Supplementary Figure S7. | DREEM-deconvoluted DMS profiles of the CR4/5 domain in HeLa cells overexpressing hTERT. |
| Supplementary Figure S8. | DREEM-deconvoluted DMS profiles of the CR4/5 domain in overexpressed WT hTR. |
| Supplementary Figure S9. | Population average DMS reactivity of the hTR t/PK domain within biochemically purified telomerase RNP complexes. |

**Supplementary Table S1. Oligonucleotides used in the study (listed 5'→3')**

| | |
|---|---|
| hTR RT, UMI, partial Nextera | GTCTCGTGGGCTCGGAGATGTGTATAAGAGACAGNNNNGCATGTGTGAGCCGAGTCCTGGGTGC |
| hTR second strand, UMI, partial Nextera | TCGTCGGCAGCGTCAGATGTGTATAAGAGACAGNNNNGGGTTGCGGAGGGTGGGCCTG |
| hTR PK amplicon, partial Nextera | TCGTCGGCAGCGTCAGATGTGTATAAGAGACAGTTCGCCCCTCCCGGGGAC |
| hTR TWJ amplicon, partial Nextera | GTCTCGTGGGCTCGGAGATGTGTATAAGAGACAGTGCCTCCGGAGAAGCCCCG |
| Forward hTR cloning (pBS-U1-hTR) | TCTAGAACTAGTGGATCCCCCGGG |
| Reverse hTR cloning (pBS-U1-hTR) | CGAGGTCGACGGTATCGATAAGCTTG |
| M2 Megaprimer | GGGTGCCACCGCAGAAGCCCCGGGCCGAC |
| M3 Megaprimer | GCCACCGCAGAAGCGCCGGGCCGACCGCGGCC |
| Nextera_Primer_A1 | AATGATACGGCGACCACCGAGATCTACACTAGATCGCTCGTCGGCAGCGTCAGATG |
| Nextera_Primer_A2 | AATGATACGGCGACCACCGAGATCTACACCTCTGTATTCGTCGGCAGCGTCAGATG |
| Nextera_Primer_A3 | AATGATACGGCGACCACCGAGATCTACACGATCAGCATCGTCGGCAGCGTCAGATG |
| Nextera_Primer_A4 | AATGATACGGCGACCACCGAGATCTACACACAGTATGTCGTCGGCAGCGTCAGATG |
| Nextera_Primer_A5 | AATGATACGGCGACCACCGAGATCTACACGACTGGAGTCGTCGGCAGCGTCAGATG |

| | |
|---|---|
| Nextera_Primer_A6 | AATGATACGGCGACCACCGAGATCTACACACTGCATATCGTCGGCAGCGTCAGATG |
| Nextera_Primer_A7 | AATGATACGGCGACCACCGAGATCTACACTTGCATGCTCGTCGGCAGCGTCAGATG |
| Nextera_Primer_A8 | AATGATACGGCGACCACCGAGATCTACACCGAATCCTTCGTCGGCAGCGTCAGATG |
| Nextera_Primer_B1 | CAAGCAGAAGACGGCATACGAGATTCGCCTTAGTCTCGTGGGCTCGGAGATGTGTAT |
| Nextera_Primer_B2 | CAAGCAGAAGACGGCATACGAGATCTAGTACGGTCTCGTGGGCTCGGAGATGTGTAT |
| Nextera_Primer_B3 | CAAGCAGAAGACGGCATACGAGATTTCTGCCTGTCTCGTGGGCTCGGAGATGTGTAT |
| Nextera_Primer_B4 | CAAGCAGAAGACGGCATACGAGATGCTCAGGAGTCTCGTGGGCTCGGAGATGTGTAT |
| Nextera_Primer_B5 | CAAGCAGAAGACGGCATACGAGATAGGAGTCCGTCTCGTGGGCTCGGAGATGTGTAT |
| Nextera_Primer_B6 | CAAGCAGAAGACGGCATACGAGATCATGCCTAGTCTCGTGGGCTCGGAGATGTGTAT |
| Nextera_Primer_B7 | CAAGCAGAAGACGGCATACGAGATGTAGAGAGGTCTCGTGGGCTCGGAGATGTGTAT |
| Nextera_Primer_B8 | CAAGCAGAAGACGGCATACGAGATCAGCCTCGGTCTCGTGGGCTCGGAGATGTGTAT |
| Northern Blot Probe hTR 130R-27 | CTT TTC CGC CCG CTG AAA GTC AGC GAG |
| Northern Blot Probe hTR 255R-28 | GCC TCC AGG CGG GGT TCG GGG GCT GGG C |
| Northern Blot Probe U1 101R-21 | GGG GAA ATC GCA GGG GTC AGC |
| Northern Blot Probe U2 153R-22 | CGA TGC GTG GAG TGG ACG GAG C |

**Supplementary Figure S1**

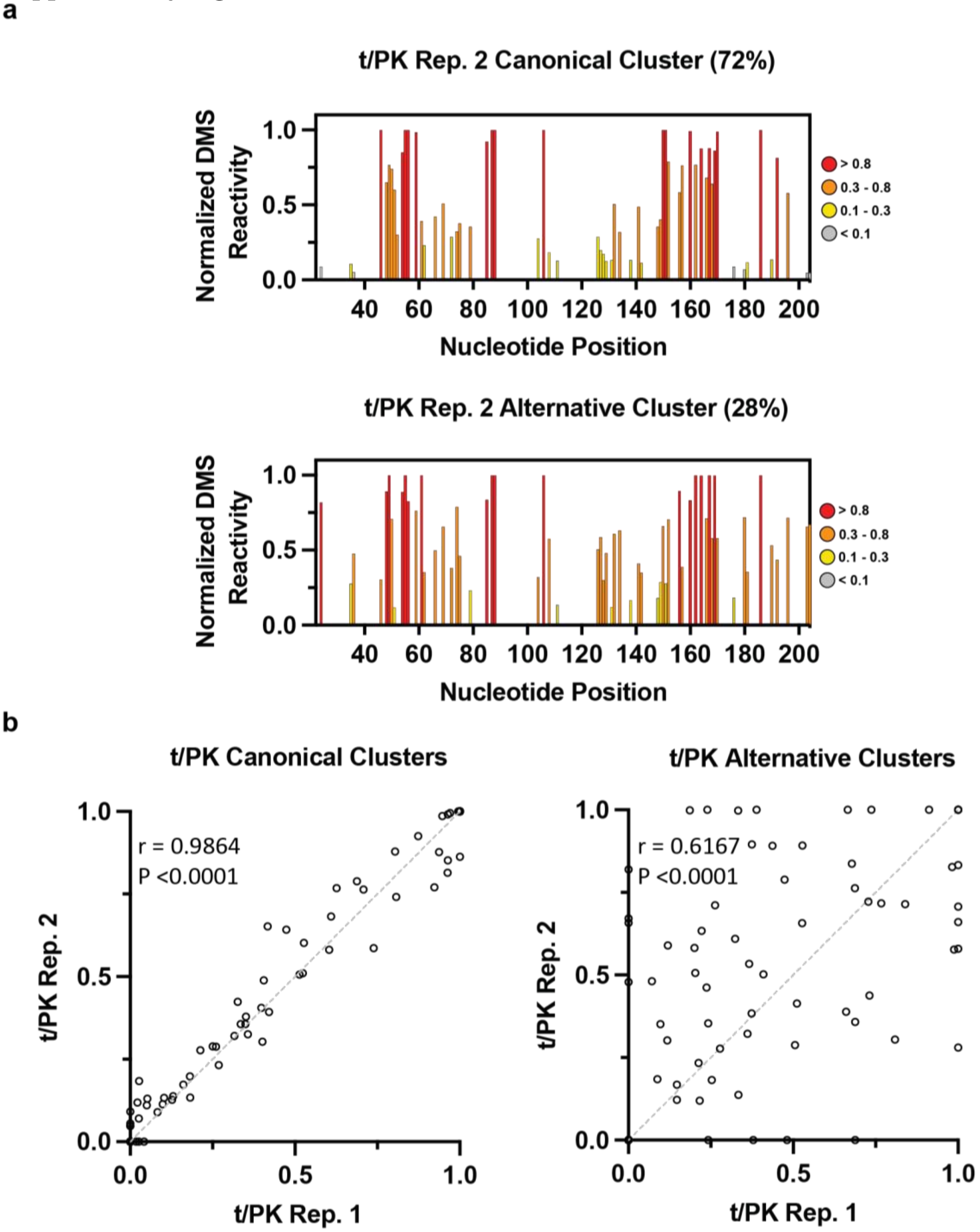


**Supplementary Figure S1. Replicate DREEM-deconvoluted DMS profiles of hTR t/PK. (a)** Normalized DMS reactivity profiles of the clusters predicted by DREEM in a biological replicate experiment. Intensity of DMS reactivity colored according to the provided legend. **(b)** Comparison of DMS reactivities from DREEM-predicted clusters of canonical (left) and alternative (right) conformations of the hTR t/PK domain. Pearson correlation (r) and P values from a two-tailed test are shown. Line of identity is shown as a grey dashed line.

**Supplementary Figure S2**

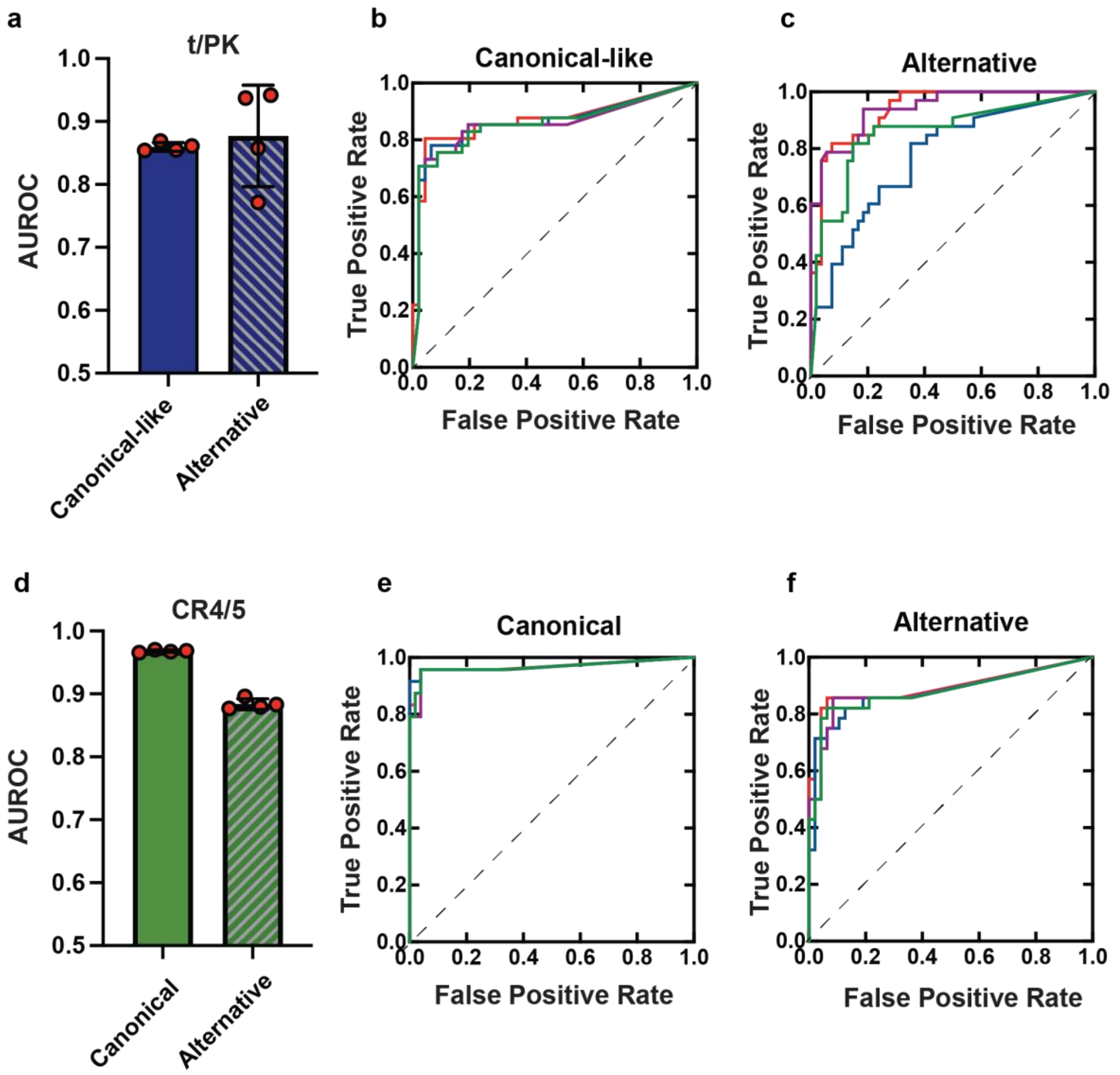


**Supplementary Figure S2. Quantitative evaluation of DMS-modification data concordance with predicted RNA secondary structure models.**

**Supplementary Figure S3**

**a**

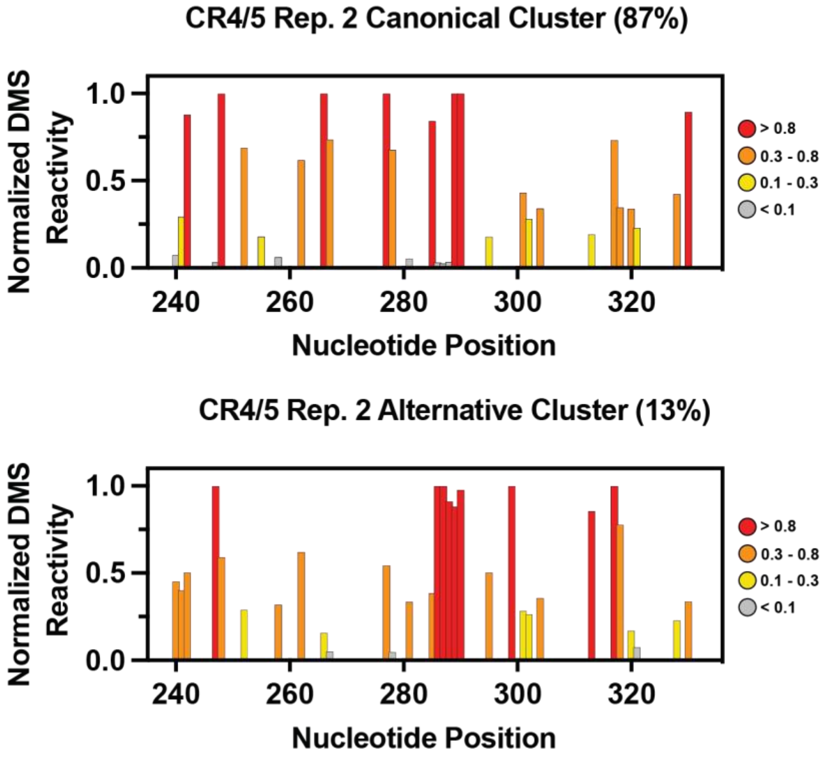


**b**

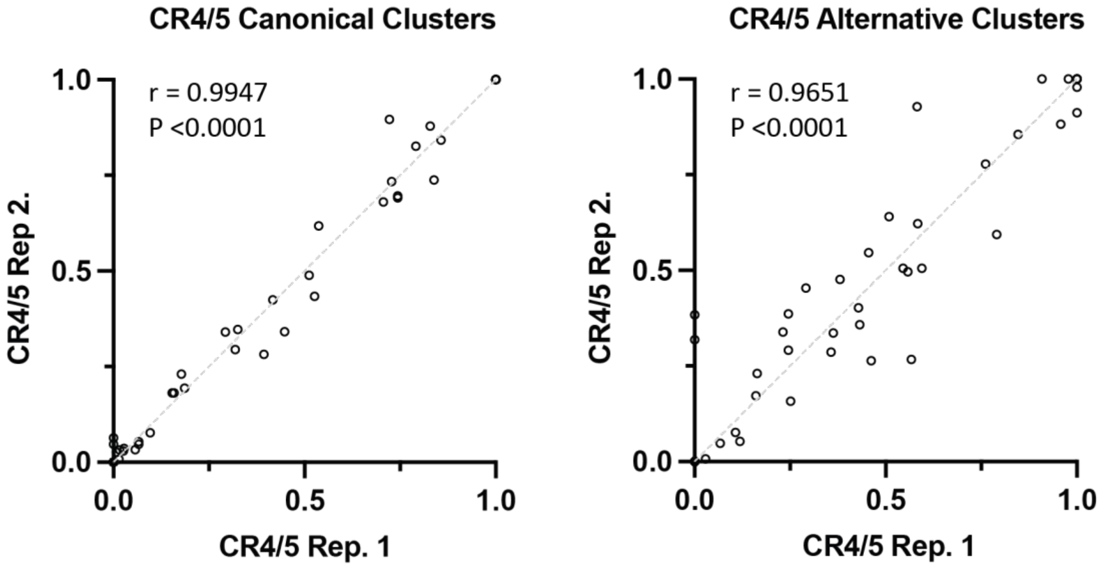


**Supplementary Figure S3. Replicate DREEM-deconvoluted DMS profiles of hTR CR4/5. (a)** Normalized DMS reactivity profiles of the clusters predicted by DREEM in a biological replicate experiment. Intensity of DMS reactivity colored according to the provided legend. **(b)** Comparison of DMS reactivities from DREEM-predicted clusters of canonical (left) and alternative (right) conformations of the hTR CR4/5 domain. Pearson correlation (r) and P values from a two-tailed test are shown. Line of identity is shown as a grey dashed line.

**Supplementary Figure S4**

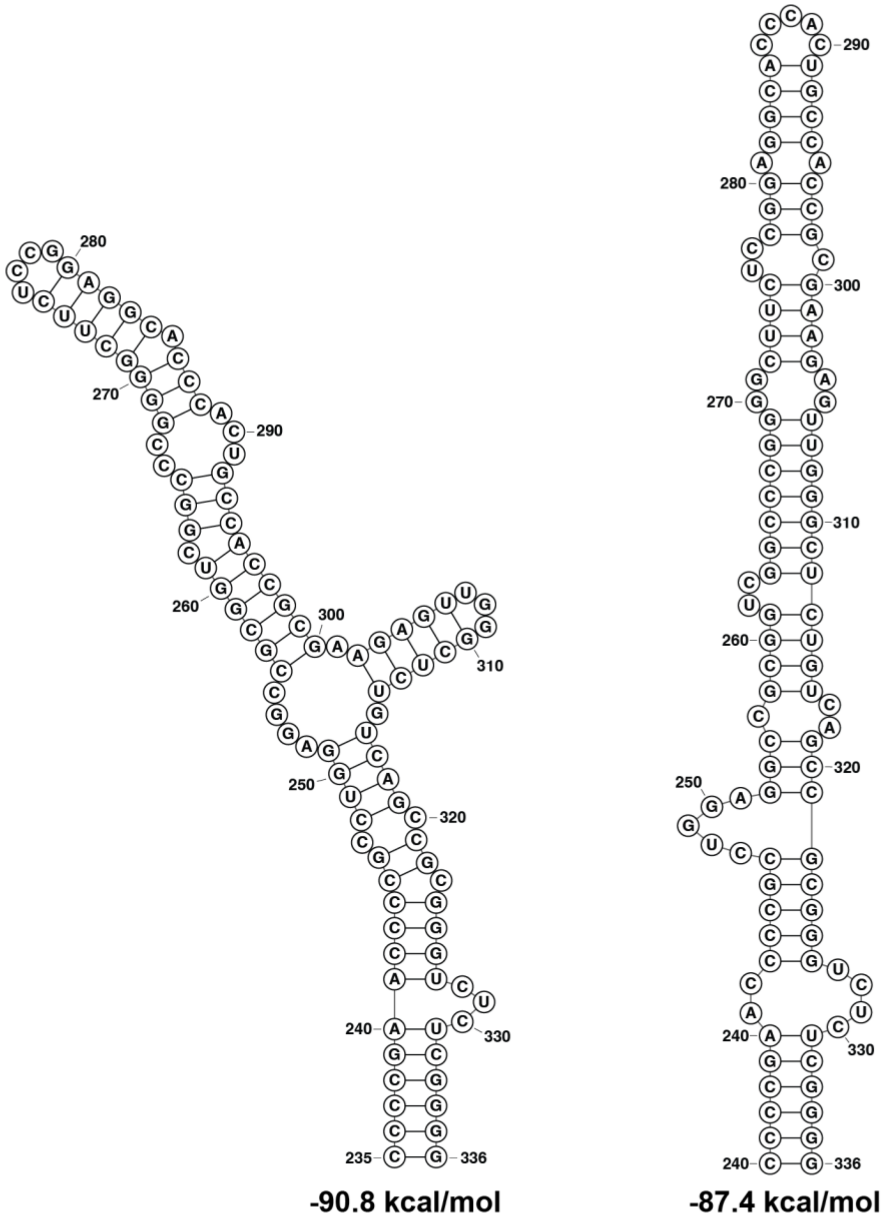


**Supplementary Figure S4. Unguided RNAstructure predictions of hTR CR4/5.** Secondary structure predictions of the hTR CR4/5 using RNAstructure default parameters and no DMS reactivity yield the canonical (left) and alternative (right) conformations with similar predicted free energies.

**Supplementary Figure S5**

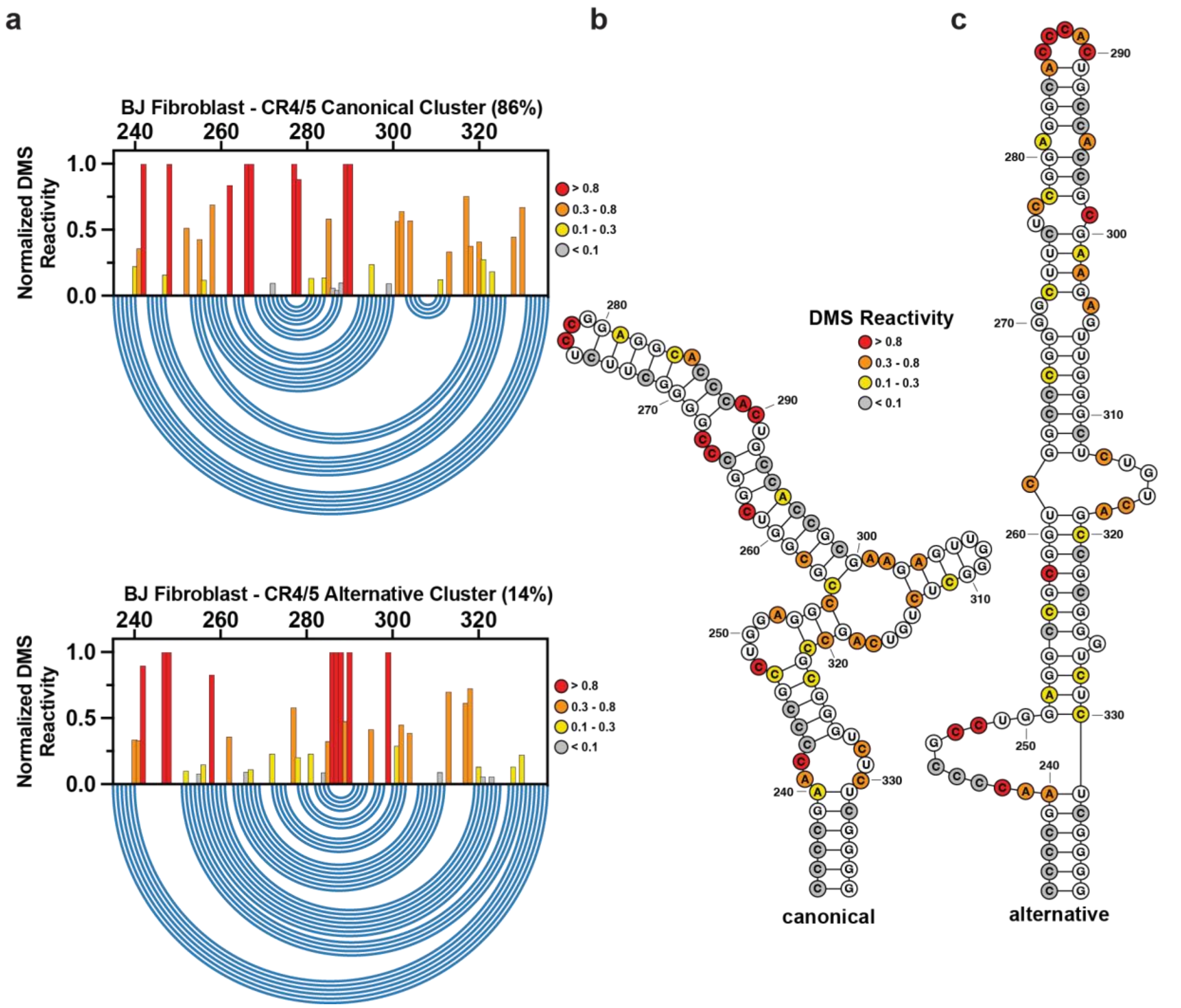


**Supplementary Figure S5. DREEM-deconvoluted DMS profiles of the CR4/5 domain in BJ Fibroblasts. (a)** Normalized DMS reactivity of the hTR CR4/5 domain in BJ fibroblast cells. Intensity of DMS reactivity according to the provided legend. Blue arcs designate the base pairing pattern of the data-guided predicted CR4/5 secondary structure. **(b)** Data-guided secondary structure prediction of the CR4/5 domain from the canonical cluster of DMS reactivities. **(c)** Data-guided secondary structure of the CR4/5 domain from the alternative cluster of DMS reactivities.

**Supplementary Figure S6**

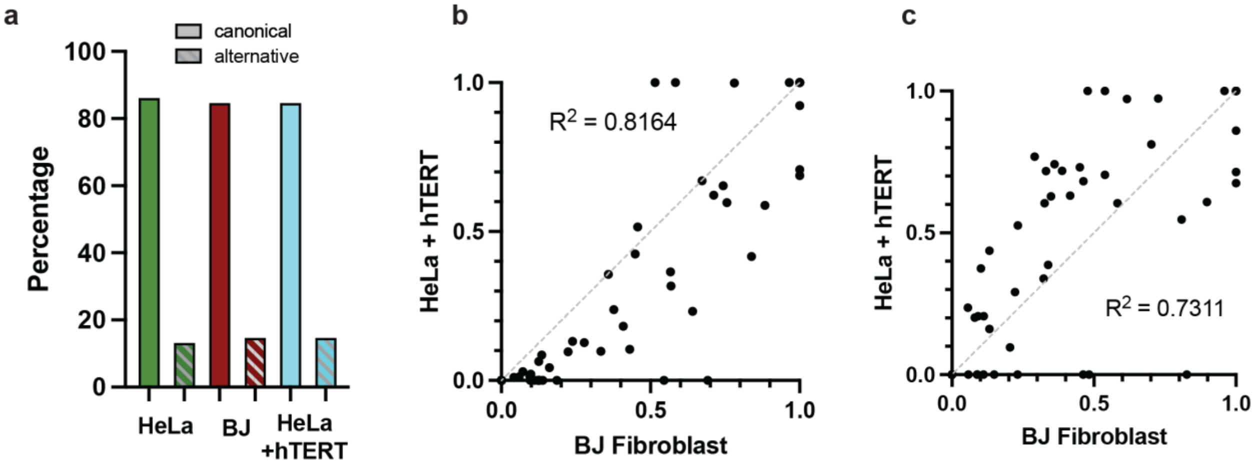


**Supplementary Figure S6. Comparison of DREEM-deconvolution results across different cell types. (a)** Bar graph depicting the proportion of the canonical conformation (solid bars) and the alternative conformation (hatched bars) of the CR4/5 domain in HeLa cells (green bars), BJ fibroblasts (red bars), and HeLa cells transiently transfected with a plasmid encoding hTERT with an N-terminal FLAG tag (light blue bars). Comparison of DMS reactivities from DREEM-predicted clusters of canonical (**b**) and alternative (**c**) conformations of the hTR CR4/5 domain between HeLa cells overexpressing hTERT (y-axis) and BJ fibroblasts (x-axis). Pearson correlation (r) and P values from a two-tailed test are shown. Line of identity is shown as a grey dashed line.

### Supplementary Figure S7

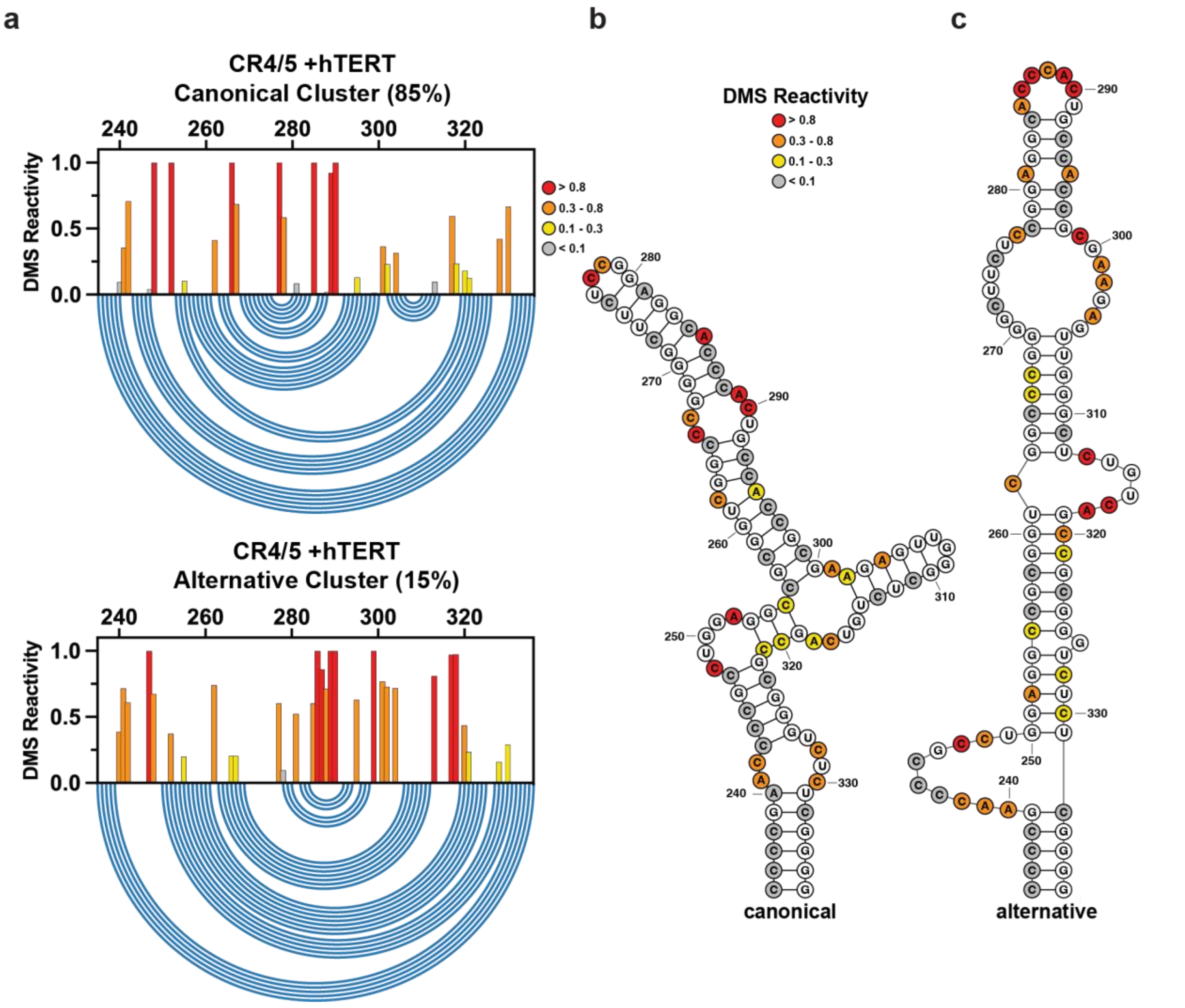


**Supplementary Figure S7. DREEM-deconvoluted DMS profiles of the CR4/5 domain in HeLa cells overexpressing hTERT. (a)** Normalized DMS reactivity of the hTR CR4/5 domain in HeLa cells transiently transfected with plasmid encoding an N-terminal tagged hTERT. Intensity of DMS reactivity according to the provided legend. Blue arcs designate the base pairing pattern of the data-guided predicted CR4/5 secondary structure. **(b)** Data-guided secondary structure prediction of the CR4/5 domain from the canonical cluster of DMS reactivities. **(c)** Data-guided secondary structure of the CR4/5 domain from the alternative cluster of DMS reactivities.

**Supplementary Figure S8**

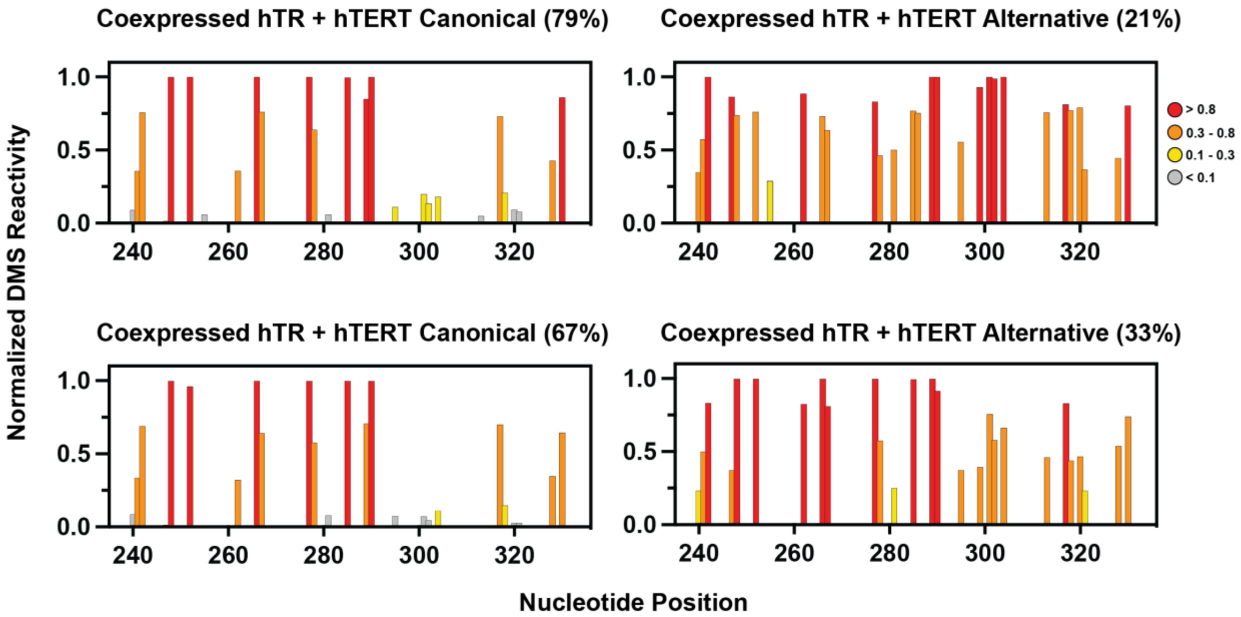


**Supplementary Figure S8. DREEM-deconvoluted DMS profiles of the CR4/5 domain in cells co-expressing WT hTR and hTERT.** DMS profiles of the two clusters predicted by DREEM from cells overexpressing WT hTR and hTERT. Cluster abundances are depicted as percentages. Results from two independent experiments are shown.

**Supplementary Figure S9**

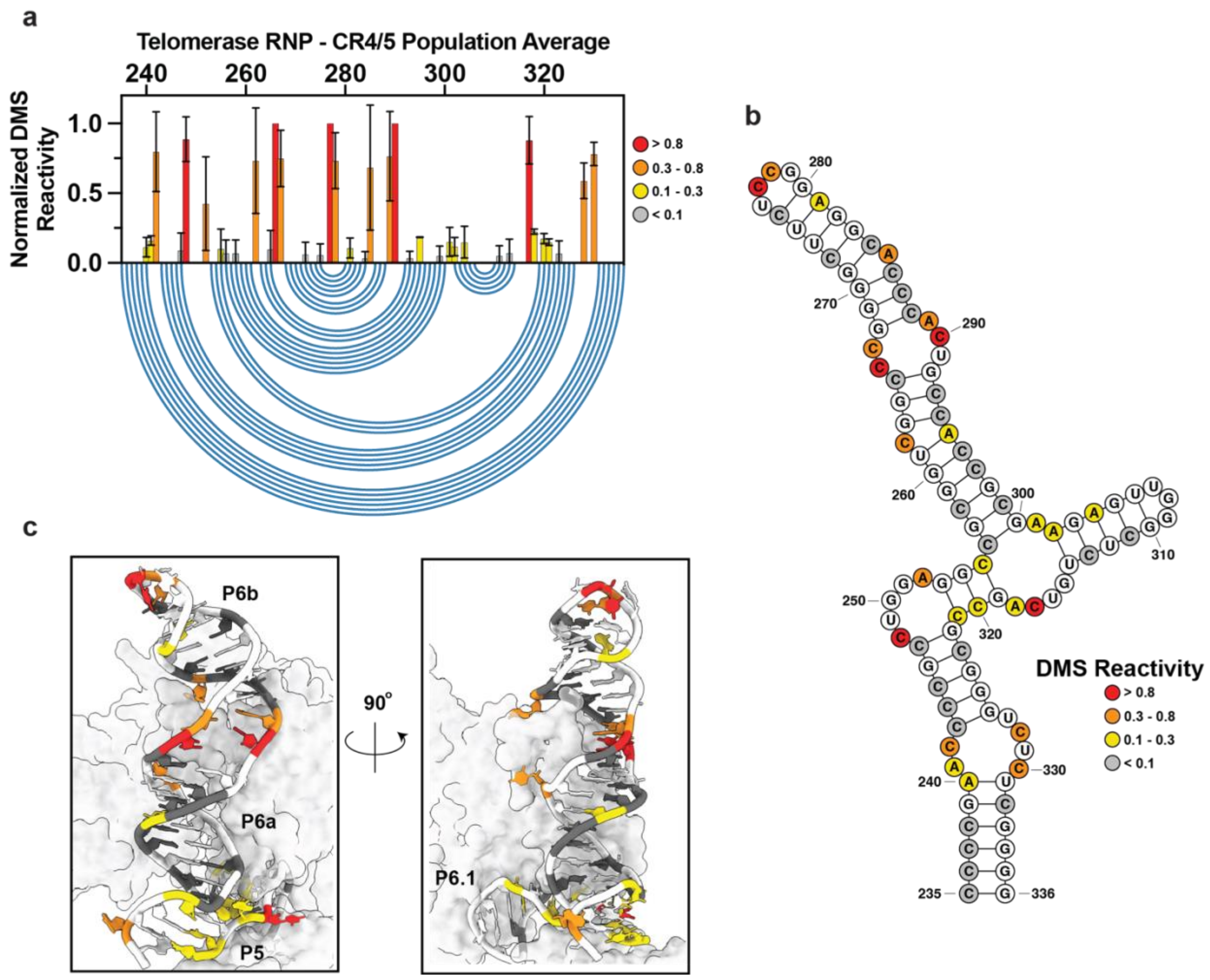


**Supplementary Figure S9. Population average DMS reactivity of the hTR CR4/5 domain within biochemically purified telomerase RNP complexes.** (**a**) Normalized DMS reactivity of hTR CR4/5 domain. Intensity of DMS reactivity colored according to the provided legend. Blue arcs designate the base pairing pattern of the data-guided predicted CR4/5 secondary structure. (**b**) Data-guided secondary structure prediction of the CR4/5 domain from the population average of DMS reactivities. **(c)** DMS reactivity overlaid onto the cryo-EM model of assembled telomerase.
